# Tamoxifen improves glucose tolerance in a delivery, sex, and strain-dependent manner in mice

**DOI:** 10.1101/467266

**Authors:** Alexis M. Ceasrine, Eugene E. Lin, David N. Lumelsky, Nelmari Ruiz-Otero, Erica D. Boehm, Rejji Kuruvilla

## Abstract

Tamoxifen, a selective estrogen receptor modulator, is widely used in mouse models to temporally control gene expression but is also known to affect body composition. Here, we report that tamoxifen has significant and sustained effects on glucose tolerance, independent of effects on insulin sensitivity, in mice. Intraperitoneal, but not oral, tamoxifen delivery improved glucose tolerance in three inbred mouse strains. The extent and persistence of tamoxifen-induced effects were sex- and strain-dependent. These findings highlight the need to revise commonly used tamoxifen-based protocols for gene manipulation in mice by including longer chase periods following injection, oral delivery, and the use of tamoxifen-treated littermate controls.

## INTRODUCTION

Genetic mouse models have been widely used for the study of human metabolic disorders, such as diabetes. The Cre/LoxP system provides a powerful tool to manipulate gene expression in a tissue- and/or temporal-specific manner in mice. Temporal control of the Cre/LoxP system is frequently accomplished with the Cre^ER^ variant, which only enters the nucleus to induce DNA recombination upon addition of the estrogen receptor modulator tamoxifen (1, 2). Despite the widespread use of Cre^ER^/LoxP mouse lines, phenotyping the effects of tamoxifen on control animals has not been well documented. Tamoxifen has been reported to significantly reduce food intake and body weight in ovariectomized rats (3). In ovariectomized mice, tamoxifen also protects against high-fat diet induced obesity and insulin resistance (4). In human breast cancer patients, tamoxifen treatment has been correlated with increased risk of diabetes (5). Together, these studies suggest that tamoxifen alters glucose homeostasis, although the effects of tamoxifen on glucose tolerance and insulin secretion have been poorly defined.

Here, we report that tamoxifen administration enhances glucose tolerance in mice. Intraperitoneal tamoxifen administration improved glucose tolerance in three commonly used mouse strains, *BALB/cJ*, *C57BL/6J*, and *129S1/SvImJ*, although the extent and persistence of enhanced glucose tolerance varied in a sex- and strain-dependent manner. In *C57BL/6J* mice, intraperitoneal tamoxifen delivery enhanced glucose-stimulated insulin secretion in males, but not, females. Tamoxifen administration had no effect on insulin sensitivity in all three mouse strains. Intriguingly, oral administration of tamoxifen through diet did not affect glucose tolerance. Together, these results highlight the need for caution in interpreting results of metabolic changes when using intraperitoneal tamoxifen delivery for temporal control of gene expression in mice, and suggest revised measures, such as extended chase periods following injection, oral delivery, and comparison to tamoxifen-treated littermate controls.

## RESULTS

### Intraperitoneal tamoxifen injections improve glucose tolerance in mice

To assess the effects of tamoxifen on glucose metabolism in mice, we employed a commonly used paradigm for tamoxifen-inducible gene inactivation in Cre^ER^/LoxP mouse models via intraperitoneal (i.p) administration of tamoxifen (100mg/kg body weight in corn oil) for 5 consecutive days (8). We assessed glucose tolerance, insulin tolerance, and glucose-stimulated insulin secretion (GSIS) *in vivo* in three inbred mouse strains: *BALB/cJ*, *C57BL/6J*, and *129S1/SvImJ*. Body weights were measured weekly and were unaffected in both sexes of all strains tested after tamoxifen injection (data not shown).

Following tamoxifen injections, we performed glucose tolerance tests on male and female mice from all three strains, 1 week after the final administration. Tamoxifen injections significantly improved glucose tolerance in male and female *BALB/cJ* and *C57BL/6J* mice (Figure 1A-F), and in female, but not male, *129S1/SvImJ* mice (Figure 1G-1I). Of note, tamoxifen-injected male *BALB/cJ* mice also showed significantly lower fasted blood glucose levels compared to controls (Figure 1A). Together, these results suggest that tamoxifen administration (i.p.) enhances glucose tolerance in mice.

**Figure 1.**
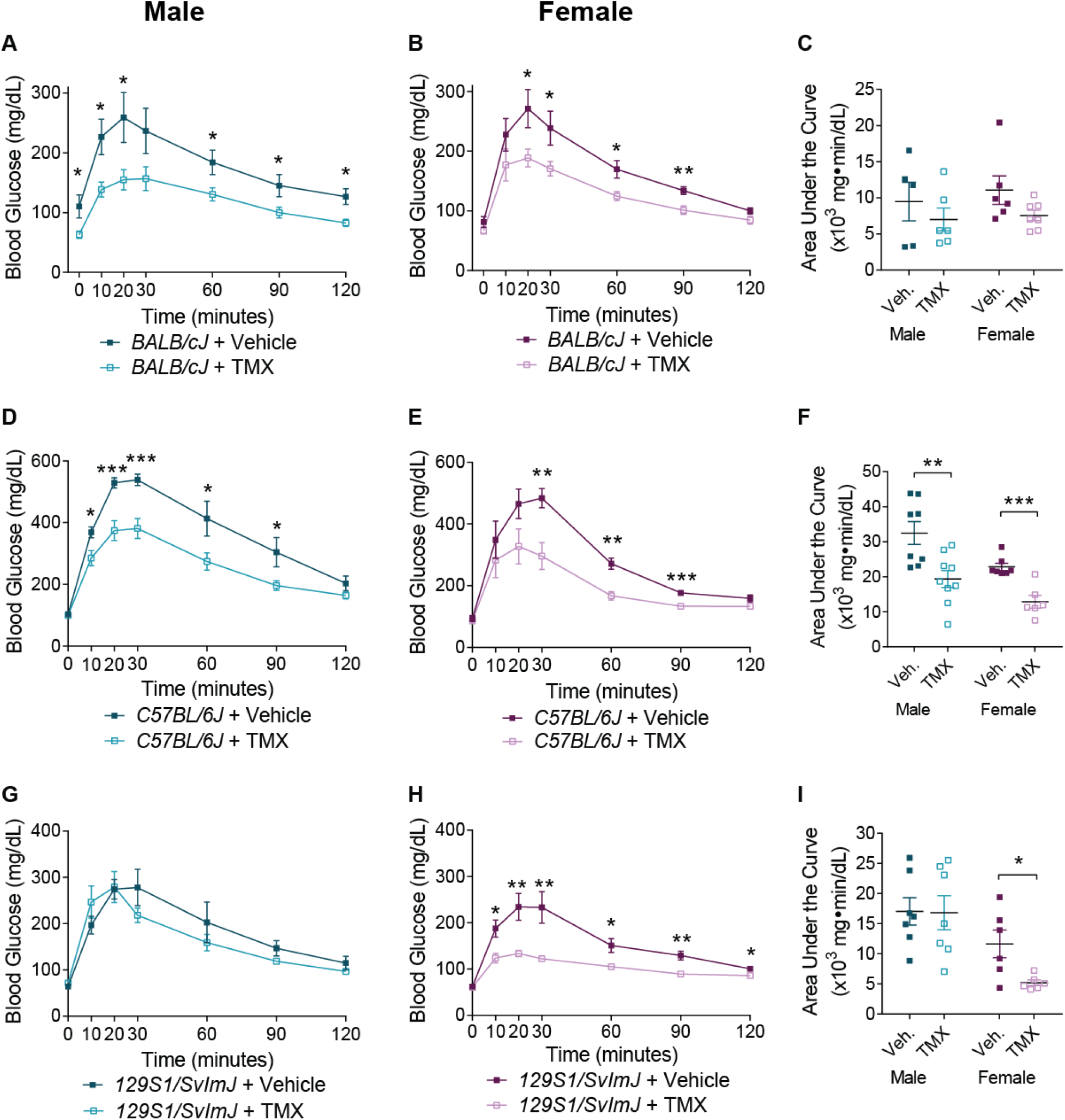
Intraperitoneal tamoxifen injections improve glucose tolerance in mice. (A) Both male and (B) female *BALB/cJ* mice show significantly improved glucose tolerance following i.p. tamoxifen (TMX) administration. Means ± SEM for n=5-7 *BALB/cJ* mice per sex per condition. (C) Area Under the Curve (AUC) for *BALB/cJ* glucose tolerance. (D) Both male and (E) female *C57BL/6J* mice show significantly improved glucose tolerance following i.p. tamoxifen administration. Means ± SEM for n= 6-9 *C57BL/6J* mice per sex per condition. (F) Area Under the Curve for *C57BL/6J* glucose tolerance. (G) Male *129S1/SvImJ* mice do not show changes in glucose tolerance in response to i.p. tamoxifen administration. Means ± SEM for n= 7 male 129S mice per condition. (H) Tamoxifen-injected female *129S1/SvImJ* mice show significantly improved glucose tolerance. Means ± SEM for n=6 female *129S1/SvImJ* mice per condition. (I) Area Under the Curve for *129S1/SvImJ* glucose tolerance. *p<0.05, **p<0.01, ***p<0.001; *t-*tests.

### Persistence of improved glucose tolerance in tamoxifen-injected mice is sex- and strain-dependent

To determine if the effects of tamoxifen injection on glucose tolerance persisted, we extended the recovery period to 21 days after the final tamoxifen injection. Three weeks following tamoxifen delivery, both male and female *BALB/cJ* mice, and male *C57BL/6J* mice still showed improved glucose tolerance (Figure 2A-D, 2F), albeit to a lower extent than the tamoxifen-induced effects observed 1 week after injections (Figure 1A-F). However, in female *C57BL/6J* mice, tamoxifen-induced enhancement of glucose tolerance was corrected by three weeks (Figure 2E-F). Female *129S1/SvImJ* mice still showed a modest improvement in glucose tolerance 3 weeks after injections (Figure 2H-I). Glucose tolerance in male *129S1/SvImJ* mice was still unaffected 3 weeks after injection (Figure 2G, 2I). Together, these results suggest that the effects of intraperitoneal tamoxifen delivery on improving glucose tolerance persist in a sex- and strain-dependent manner at 3 weeks following injections, with male mice tending to exhibit more long-lasting effects. Overall, the effects of tamoxifen on enhancing glucose tolerance 3 weeks post-injection were modest compared to 1-week post-injection, suggesting that extended recovery periods following tamoxifen delivery in Cre^ER^ mouse lines may be sufficient to avoid tamoxifen-induced effects on glucose tolerance.

**Figure 2.**
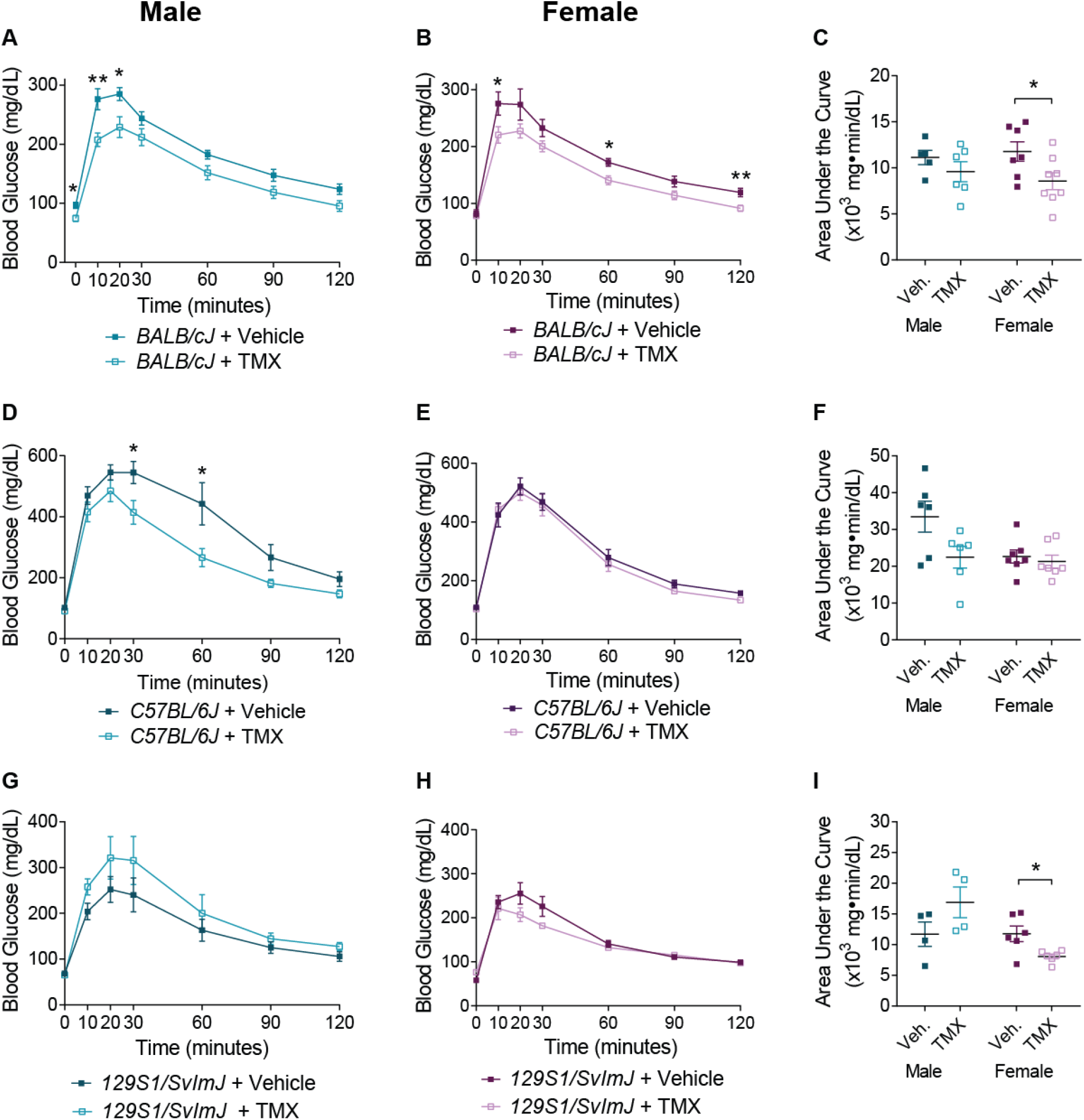
Persistence of improved glucose tolerance in tamoxifen-injected mice is sex- and strain-dependent. (A) Both male and (B) female *BALB/cJ* mice retain significantly improved glucose tolerance three weeks post-i.p. tamoxifen administration. Means ± SEM for n=5-8 *BALB/cJ* mice per sex per condition. (C) Area Under the Curve (AUC) for *BALB/cJ* glucose tolerance. (D) Male, but not (E) female, *C57BL/6J* mice retain significantly improved glucose tolerance three weeks following i.p. tamoxifen administration. Means ± SEM for n= 6-7 *C57BL/6J* mice per sex per condition. (F) Area Under the Curve for *C57BL/6J* glucose tolerance. (G) Male *129S1/SvImJ* mice do not show changes in glucose tolerance in response to i.p. tamoxifen administration. Means ± SEM for n= 4 male *129S1/SvImJ* mice per condition. (H) Glucose tolerance in tamoxifen-injected female *129S1/SvImJ* mice is modestly improved three weeks after final injection. Means ± SEM for n=6 female *129S1/SvImJ* mice per condition. (I) Area Under the Curve remains significantly lower in female *129S1/SvImJ* mice. *p<0.05, **p<0.01; *t*-tests.

### Intraperitoneal tamoxifen administration enhances GSIS in male *C57BL/6J* mice, but insulin sensitivity is unaffected in all 3 strains

Improved glucose tolerance can be caused by a variety of factors, including enhanced insulin secretion from pancreatic islets or improved insulin sensitivity in peripheral tissues.Estrogen receptor signaling has been shown to promote GSIS in isolated islets, and tamoxifen is capable of binding to pancreatic estrogen receptors (9; 10). Furthermore, tamoxifen-induced Cre-mediated recombination has been shown to persist up to 4 weeks after tamoxifen injection in pancreatic islets (11). Therefore, we asked if tamoxifen injection improves glucose tolerance by influencing insulin secretion. We assessed GSIS *in vivo* in male and female *C57BL/6J* mice, the most commonly used inbred mouse strain (12), 1 week after final intraperitoneal tamoxifen injection. Tamoxifen injections enhanced GSIS in male, but not female, *C57BL/6J* mice (Figure 3A-B), suggesting sex-specific effects of tamoxifen on insulin secretion. Insulin tolerance tests revealed that tamoxifen injections had no effect on insulin sensitivity in male and female *C57BL/6J* mice (Figure 3C-D). We also assessed insulin sensitivity in all 3 mouse strains using HOMA-IR measurements. HOMA-IR scores are estimated using fasting insulin and blood glucose levels, and a HOMA-IR score of 1 indicates optimal insulin sensitivity. This output has been adapted and verified in rodent models (13; 14). Tamoxifen injections had no effect on HOMA-IR scores in all three mouse strains, although *C57BL/6J* mice had higher HOMA-IR scores, suggesting diminished insulin sensitivity, compared to *129S1/SvImJ* and *BALB/cJ* mice (Figure 3E), consistent with previous studies (15). Together, these results suggest that tamoxifen-induced improvement of glucose tolerance in male *C57BL/6J* mice likely stems from enhanced insulin secretion. Further, enhanced insulin sensitivity is not a contributing factor to the tamoxifen-induced improved glucose tolerance in all mouse strains.

**Figure 3.**
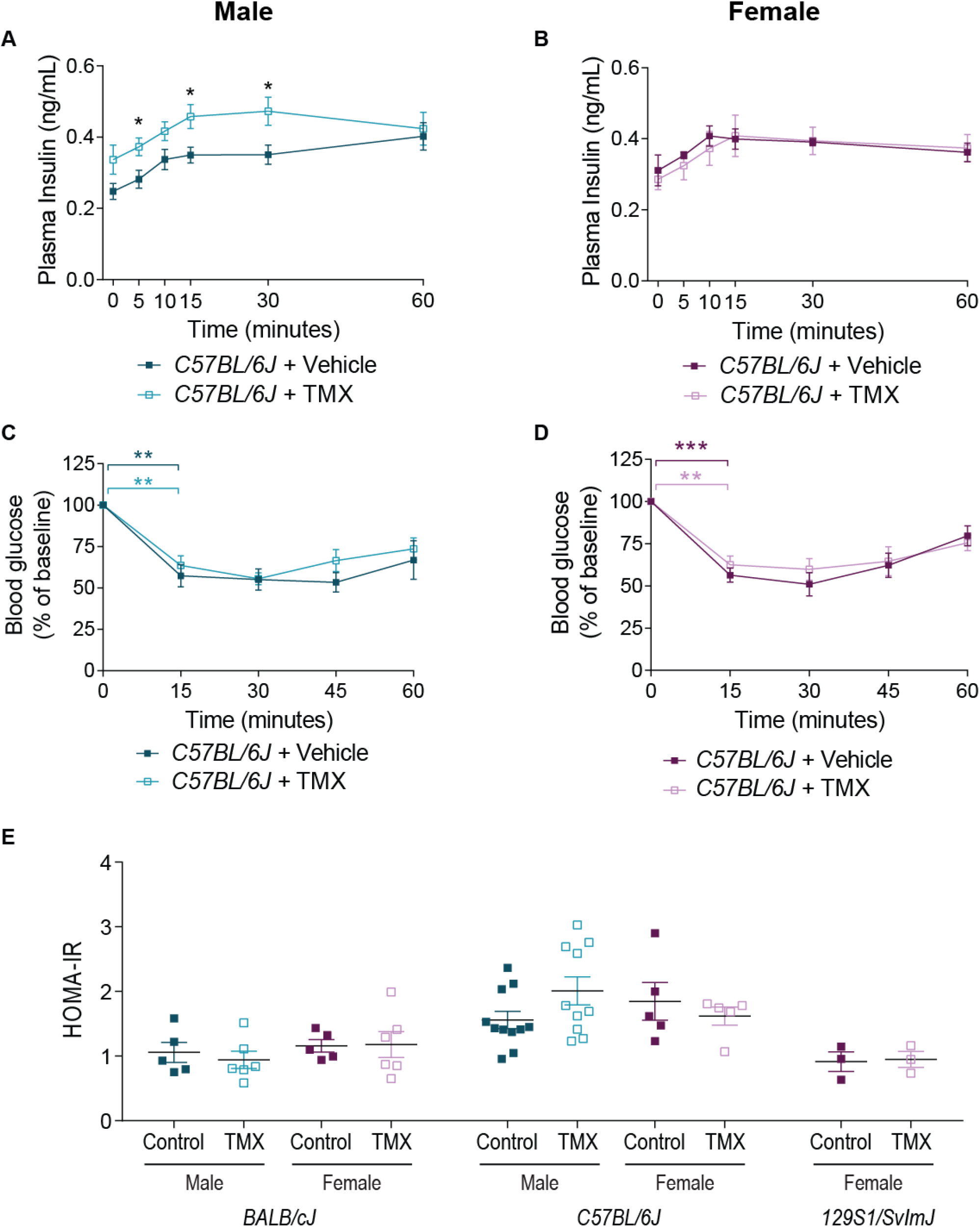
Intraperitoneal tamoxifen administration enhances GSIS in male *C57BL/6J* mice, but insulin sensitivity is unaffected in all 3 strains. (A) Glucose-stimulated insulin secretion is improved following i.p. tamoxifen administration in male *C57BL/6J* mice. Means ± SEM for n=5-11 mice per condition. *p<0.05; *t*-tests. (B) Glucose-stimulated insulin secretion is unchanged by i.p. tamoxifen administration in female *C57BL/6J* mice. Means ± SEM for n=5-8 mice per condition. (C) Insulin sensitivity is unaffected by i.p. tamoxifen administration in both male and (D) female *C57BL/6J* mice. Means ± SEM for n=5-6 *C57BL/6J* mice per condition per sex. **p<0.01, ***p<0.001; one-sample *t*-tests. (E) HOMA-IR scores for male and female *BALB/cJ* and *C57BL/6J* mice, and female *129S1/SvImJ* mice show no tamoxifen-induced changes in insulin sensitivity. Means ± SEM for n=3-11 mice per strain per condition per sex.

### Oral tamoxifen delivery does not affect glucose tolerance

Recent studies have described the use of tamoxifen-supplemented chow as an effective and non-invasive means for spatiotemporal gene activation in Cre^ER^ mouse lines (16; 17). Further, plasma tamoxifen levels are lower following oral tamoxifen delivery *versus* injection (18). Thus, we asked if oral tamoxifen delivery would also affect glucose tolerance in mice. We placed mice on commercially available tamoxifen-supplemented or control diets for 10 days with a 2-day break after the 5^th^ day. This paradigm was based on the assumption that mice consume an average of 3-4g food/day and weigh 20-25g, corresponding to approximately 40mg tamoxifen/kg body weight/day (19). Approximate tamoxifen consumption was estimated by weighing tamoxifen-supplemented chow before and after the treatment period and calculating mg tamoxifen consumed per animal (based on 250mg tamoxifen/kg of food). These analyses suggested that comparable amounts of tamoxifen were delivered between intraperitoneal injection and diet delivery (approximately 500 mg/kg total from injection versus 440 ± 77 mg/kg for males and 432 ± 73 mg/kg for females from diet (values are mean ± SD for n=10 males and 11 females from all three strains). Interestingly, glucose tolerance was unaffected by oral tamoxifen administration in males and females in all three mouse strains tested (Figures 4 A-I). These results suggest that the mode of tamoxifen administration significantly impacts its effects on glycemic control.

**Figure 4.**
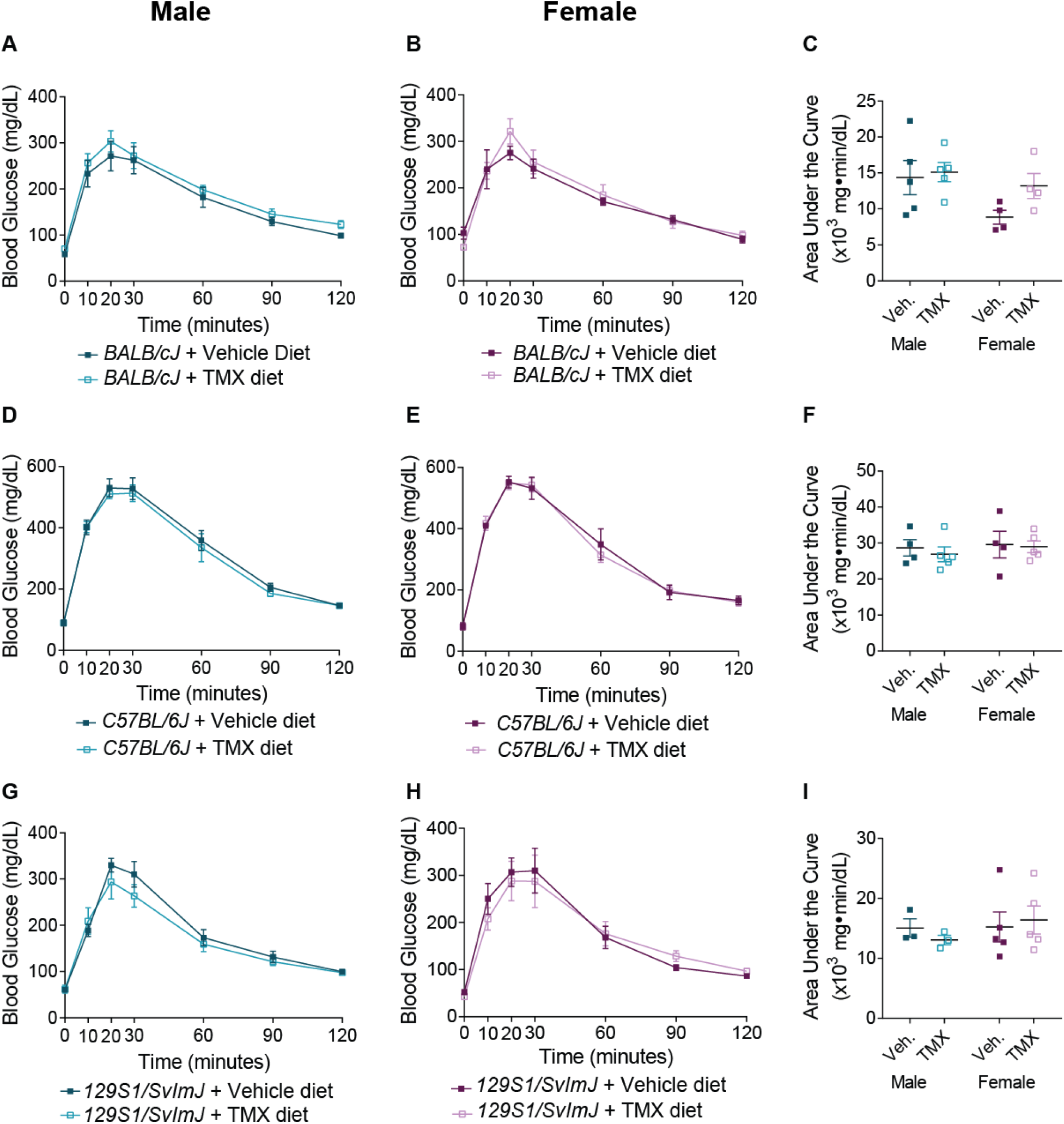
Oral tamoxifen administration does not affect glucose tolerance. (A) Male and (B) female *BALB/cJ* mice do not show changes in glucose tolerance in response to tamoxifen diet. Means ± SEM for n=4-5 *BALB/cJ* mice per sex per condition. (C) Area Under the Curve (AUC) for *BALB/cJ* glucose tolerance. (D) Male and (E) female *C57BL/6J* mice do not show changes in glucose tolerance in response to tamoxifen diet. Means ± SEM for n=4-5 *C57BL/6J* per sex per condition. (F) Area Under the Curve for *C57BL/6J* glucose tolerance. (G) Male and (H) female *129S1/SvImJ* mice do not show changes in glucose tolerance in response to tamoxifen diet. Means ± SEM for n=3-5 *129S1/SvImJ* mice per sex per condition. (I) Area Under the Curve for *129S1/SvImJ* glucose tolerance.

## DISCUSSION

Here, we report that intraperitoneal tamoxifen administration to three common inbred mouse strains induced a significant, and often sustained, improvement of glucose tolerance without affecting insulin sensitivity. Interestingly, male, but not female, *C57BL/6J* mice showed increased glucose-stimulated insulin secretion *in vivo*. Estrogen signaling has been shown to promote neural-dependent insulin secretion in males (20), as well as promote insulin secretion from isolated islets (10), and either of these effects might explain the tamoxifen-dependent insulin secretion in male *C57BL/6J* mice. Enhanced glucose tolerance in female *C57BL/6J* mice could stem from insulin-independent glucose uptake mechanisms. Previous studies suggest that insulin-independent glucose uptake is prominent during hyperglycemia in tissues such as skeletal muscle (21). Notably, estrogen receptor signaling in skeletal muscle promotes surface localization of the glucose transporter, GLUT4, in an insulin-independent manner (22). These findings, together with our observations, suggest that tamoxifen may regulate glucose tolerance in female *C57BL/6J* mice by promoting insulin-independent glucose uptake.

Previous studies have reported on sex and strain differences in glucose metabolism in mice (15). Our results indicate that the magnitude and persistence of tamoxifen-induced enhancement of glucose tolerance were sex- and strain-dependent. The persistence of tamoxifen in the body has been reported to be age-dependent, with older animals being unable to effectively clear tamoxifen (23). We used young animals (2 months of age), and tamoxifen-injected animals were always compared to age-matched vehicle-injected controls. Therefore, age is unlikely to be a factor in the observed sex-and strain-dependent effects of tamoxifen on glucose tolerance. Tamoxifen has also been shown to persist differentially in different tissues. For example, tamoxifen has been shown to have acute effects in the brain (23), and more long-lasting effects in pancreatic islets with tamoxifen-induced Cre activity persisting for as long as 4 weeks following i.p. injections (11). Therefore, tamoxifen may act differently in distinct tissues in a sex- and strain-specific manner to influence glucose tolerance.

Intriguingly, we found that oral administration of tamoxifen did not affect glucose tolerance. Compared to injection-based tamoxifen administration, oral tamoxifen delivery results in lower plasma tamoxifen levels but higher circulating levels of tamoxifen metabolites (18). These findings suggest that enhanced glucose tolerance seen following i.p. tamoxifen administration is likely due to tamoxifen itself, and not one of its metabolites. Improved glucose tolerance with tamoxifen injection might be due to the higher and persistent circulating tamoxifen levels compared to oral administration. Together, our findings highlight the need to revise commonly used tamoxifen-based protocols for gene manipulation in mice, including considering oral delivery, allowing longer chase periods following injection, and the use of tamoxifen-treated littermate controls.

## MATERIALS AND METHODS

### Animals

All procedures relating to animal care and treatment conformed to The Johns Hopkins University Animal Care and Use Committee (ACUC) and NIH guidelines. Animals were group housed in a standard 12:12 light-dark cycle. Animals of both sexes were used for all analyses, and the sex of the animals is clearly noted throughout. *BALB/cJ* (stock # 000651), *C57BL/6J* (stock # 000664), and *129S1/SvImJ* (stock #002448) mice were obtained from Jackson Laboratories.

### Tamoxifen Administration

#### Intraperitoneal Injection

At 6 weeks of age, mice were injected intraperitoneally with 100 mg/kg tamoxifen in corn oil, or corn oil alone, daily for five days. Metabolic assays were performed 1 to 3 weeks after the final injection.

#### Oral administration

Beginning at 5 weeks of age, mice were switched from standard chow to either control diet (Envigo TD.07570) or tamoxifen diet (Envigo TD.130856) for 10 days with a 2-day break on standard chow after the 5^th^ day. Since oral tamoxifen administration has been reported to reduce food intake and body weight (6), mice were given 6 days to recover on standard chow before glucose tolerance testing.

### Glucose tolerance and *in vivo* insulin secretion

Mice were individually housed and fasted overnight (16 hours) before intraperitoneal glucose injection (2g/kg glucose tolerance or 3g/kg insulin secretion). Blood glucose levels were measured with a OneTouch Ultra 2 glucometer and plasma insulin levels were measured with an Ultrasensitive Insulin ELISA (Crystal Chem). Area Under the Curve (AUC) was determined for glucose tolerance for each animal using GraphPad Prism 7.

### Insulin tolerance

Mice were individually housed with food overnight before 0.75U/kg intraperitoneal insulin injection (Novolin-R, Novo Nordisk). Blood glucose levels were measured with a OneTouch Ultra 2 glucometer.

### HOMA-IR calculations

Mice were individually housed and fasted overnight (16 hours). Fasting plasma insulin and blood glucose levels were used to calculate the HOMA-IR score as: (fasting plasma insulin (mmol/L) * blood glucose (mmol/L))/22.5 (7).

## ACKNOWLEDGEMENTS

This work was supported by a NIH grant (DK108267) to R.K., NRSA post-doctoral fellowship (F32DK116482) to E.L. A.C., N.R-O., and E.B. are all supported by a NIH training grant (T32GM007231) awarded to the JHU CMDB program. A.C. designed and performed experiments and data analysis and wrote the manuscript. E.L. performed experiments and reviewed and edited the manuscript. D.L., N.R-O., and E.B. performed experiments. R.K. designed experiments and reviewed and edited the manuscript. A.C. is the guarantor of this work and, as such, had full access to all the data in the study and takes responsibility for the integrity of the data and the accuracy of data analysis.

